# Hierarchy of transcriptomic specialization across human cortex captured by myelin map topography

**DOI:** 10.1101/199703

**Authors:** Joshua B. Burt, Murat Demirtaş, William J. Eckner, Natasha M. Navejar, Jie Lisa Ji, William J. Martin, Alberto Bernacchia, Alan Anticevic, John D. Murray

## Abstract

Hierarchy provides a unifying principle for the macroscale organization of anatomical and functional properties across primate cortex, yet the microscale bases of specialization across human cortex are poorly understood. Cortical hierarchy is conventionally informed by invasive measurements of long-range projections, creating the need for a principled proxy measure of hierarchy in humans. Moreover, cortex exhibits marked interareal variation in patterns of gene expression, yet organizing principles of its transcriptional architecture remain unclear. We hypothesized that functional specialization of human cortical microcircuitry involves hierarchical gradients of gene expression. We found that a noninvasive neuroimaging measure, the MRI-derived myelin map, reliably indexes hierarchy and closely resembles the dominant pattern of transcriptomic variation across human cortex. We found strong hierarchical gradients in expression profiles of genes related to microcircuit function and neuropsychiatric disorders. Our findings suggest that hierarchy defines an axis shared by the transcriptomic and anatomical architectures of human cortex, and that hierarchical gradients of microscale properties contribute to macroscale specialization of cortical function.

---

The neocortex of human and nonhuman primates exhibits interareal patterns of structural and functional variation. Cortical areas are distinguished by differences in their cellular composition, laminar differentiation, and long-range anatomical connectivity. Primate cortex also is also characterized by large-scale gradients of specialization in physiology and function, including in representational selectivity^1–3^ and dynamics of intrinsic activity^4,5^. Yet it remains unclear how the large-scale functional architecture of cortex may be subserved by specialization of local microcircuitry.

Recently, analysis of the molecular composition of cortical microcircuitry has been revolutionized by advances in large-scale high-throughput transcriptomics, which can produce genome-wide maps of gene expression levels across brain areas. Datasets such as the Allen Human Brain Atlas (AHBA) have revealed a rich transcriptomic architecture characterized by spatially heterogeneous gene expression profiles across areas of the human brain^6–8^. Interareal transcriptional diversity has been related to differences in cortical function, including the spatiotemporal structure of intrinsic network activity^7,9–11^, and to spatially heterogeneous patterns of anatomical connectivity^11,12^. However, unifying principles for the macroscale organization of structural, functional, and transcriptional differences across human cortex are still unknown.

A parsimonious principle for the large-scale anatomical and functional organization of nonhuman primate cortex is the concept of cortical hierarchy^2–4,13–16^. Anatomical hierarchy, defined as a globally self-consistent ordering of cortical areas according to characteristic laminar patterns of interareal projections, has been studied extensively in monkeys through histological tract-tracing methods^13–15^. The ordering of cortical areas along the anatomical hierarchy, which situates early sensory areas toward the bottom and higher-order association areas toward the top of the hierarchy, has also been found to align with their functional organization in sensory processing hierarchies^13,15^. We hypothesized that the transcriptomic architecture of human cortex is also hierarchically organized, such that the functional specialization of human cortical microcircuitry involves hierarchical gradients of gene expression. However, the highly invasive nature of the tract-tracing data acquisition procedures which are required to index hierarchy in nonhuman primates has thus far precluded analogous investigations of cortical organization in humans, thereby creating the need for noninvasive alternative measures.

To address these open questions, we analyzed transcriptomic, anatomical, and neuroimaging data from humans and monkeys to study the hierarchical organization of microcircuit specialization across human cortex. We found that a noninvasive structural neuroimaging measure, the MRI-derived myelin map^17^, provides a proxy for anatomical hierarchy in primate cortex. To test for hierarchical gradients in gene expression, we then compared the spatial expression profiles of genes in the AHBA to the topography of the human myelin map. We found strong hierarchical gradients in expression profiles of genes related to synaptic physiology, cell-type specificity, and cortical cytoarchitecture, in line with anatomical measurements in monkey. Furthermore, we observed a remarkably close correspondence between myelin map topography and the dominant spatial pattern of gene expression variation across human cortex. Finally, we found that hierarchically patterned genes are preferentially associated with functional processes and brain disorders. Overall, these findings suggest that hierarchy defines an axis shared by the transcriptomic and anatomical architectures of human cortex, and that hierarchical gradients of microscale properties shape the macroscale specialization of cortical function.

## Results

### Myelin maps noninvasively capture anatomical hierarchy

To enable the study of hierarchy in human cortex, we first sought to establish a noninvasive neuroimaging measure that can serve as a proxy measure for indexing anatomical hierarchy. One measure we examined was the cortical myelin map, a structural neuroimaging map which can be measured as the contrast ratio of T1- to T2-weighted (T1w/T2w) magnetic resonance images^17^. The MRI-derived myelin map provides a noninvasive *in vivo* correlate of gray-matter intracortical myelin content and captures established anatomical borders between cytoarchitecturally delineated cortical areas^17^. Motivated by the empirical observation that myelin map values are high in primary sensory cortex (visual, somatosensory, auditory) and low in association cortex, homologously in human and monkey (Fig. 1a–c, Extended Data Fig. 1), we hypothesized that the cortical myelin map provides a noninvasive proxy for areas’ hierarchical positions through this inverse myelin-hierarchy relationship.

**Figure 1:**
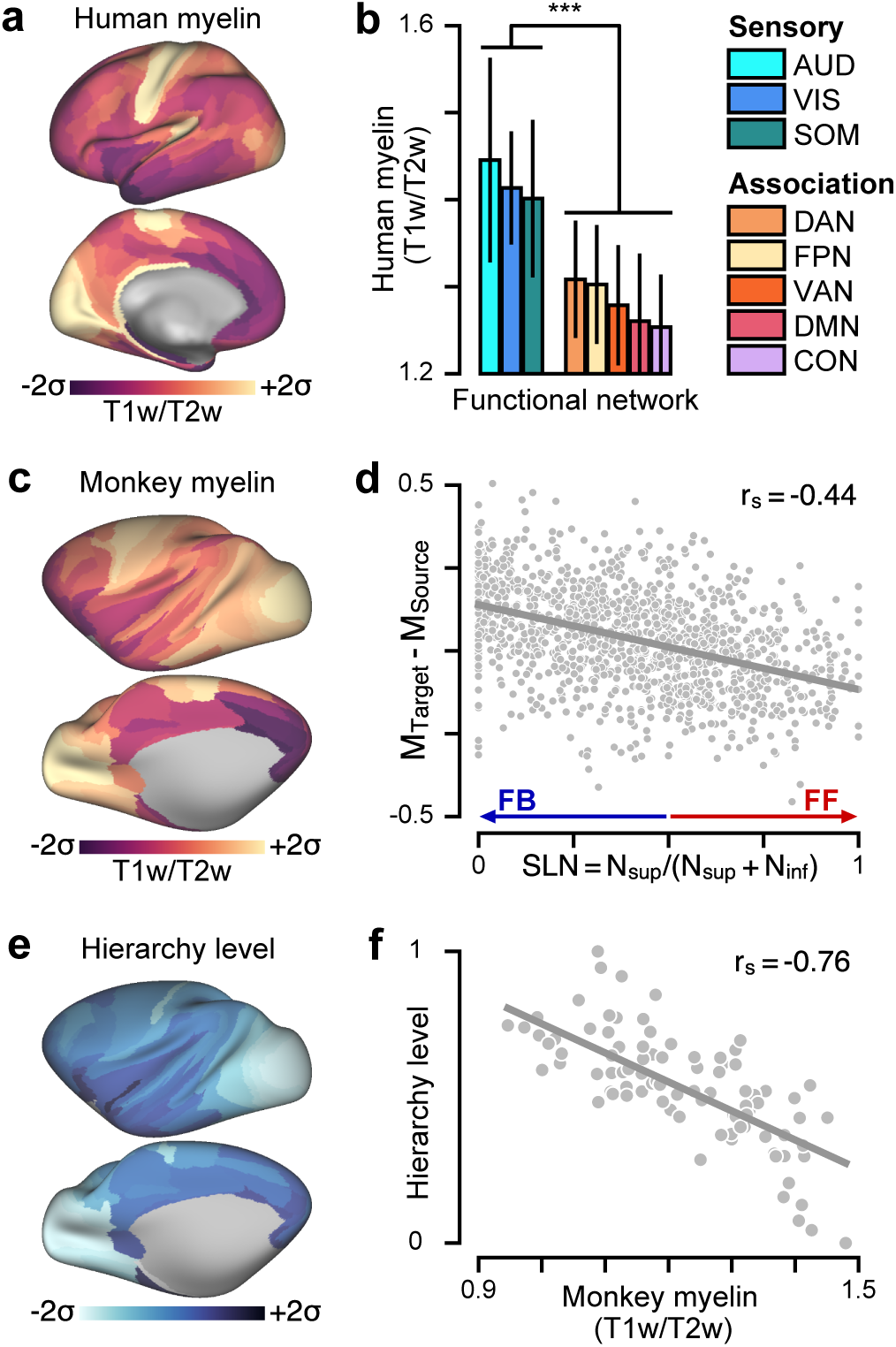
Cortical myelin map topography noninvasively captures the hierarchical organization of primate cortex. **(a)** The parcellated human cortical myelin map (T1w/T2w MRI signal) exhibits high values in primary sensory cortical areas relative to association areas. **(b)** Human myelin map values are significantly lower in functionally defined association networks than in sensory networks (*P <* 10^*−*3^; Wilcoxon signed-rank test) (Fig. 1c). Error bars mark the std. dev. across areas. **(c)** The parcellated macaque monkey myelin map topography is similar to that of the human. **(d)** Myelin map variation predicts feedforward (FF) and feedback (FB) interareal projections in monkey cortex, as quantified by the fraction of labeled supragranular layer neurons (SLN) in the source area. High and low SLN correspond to FF and FB projection motifs, respectively. SLN significantly correlates with the difference in myelin map values between target and source areas (*τ*_*s*_ = −0.44, *P <* 10^*−*5^; Spearman rank correlation). **(e)** Hierarchy levels across cortical areas are estimated by fitting a generalized linear model to predict SLN from pairwise hierarchical distance. **(f)** Hierarchy levels are reliably predicted by the myelin map values in monkey cortex (*τ*_*s*_ = *−*0.76, *P <* 10^*−*5^).

We validated the myelin map as a proxy measure of anatomical hierarchy in monkey cortex by comparing myelin map values to the hierarchy levels derived from conventional tract-tracing approaches which quantify long-range interareal projections and their laminar specificity^15^. These laminar connectivity data are used to specify a globally optimal hierarchical ordering of cortical areas, such that lower areas send feedforward projections to higher areas, and higher areas send feedback projections to lower areas^13–15,18^ (Extended Data Fig. 2). Feedforward and feedback projections primarily originate from the supragranular and infragranular cortical layers, respectively^13,15^. At the level of individual projections, we found that the difference in myelin map values between connected areas is correlated with the laminar feedforward/feedback structure of the connection (Fig. 1d). Globally, we found a strong negative correlation between anatomical hierarchy and myelin map value (*r*_*s*_ = *−*0.76, *P <* 10^*−*5^; Spearman rank correlation) (Fig. 1e,f).

**Figure 2:**
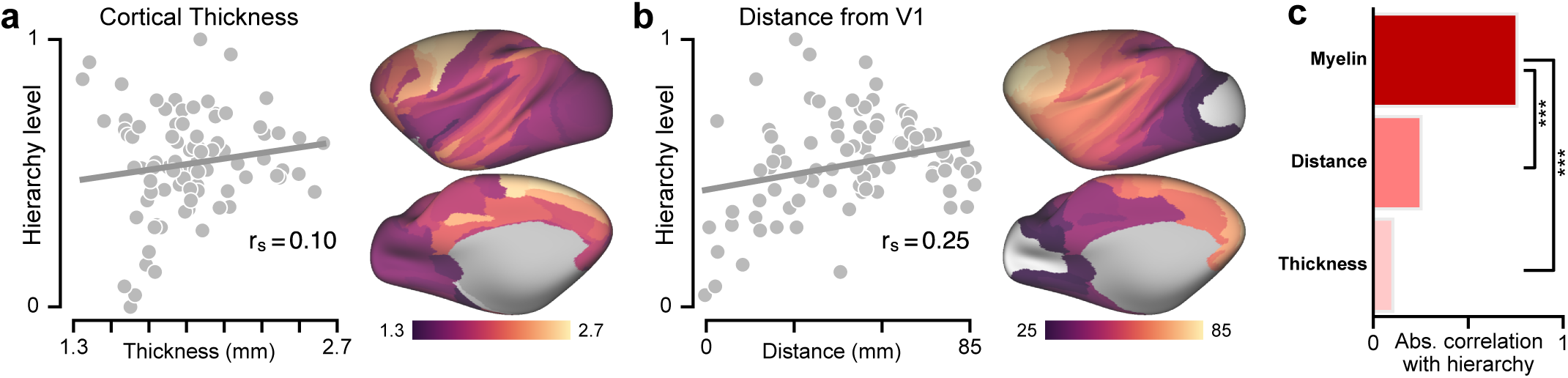
Hierarchy in monkey cortex is better captured by the myelin map (T1w/T2w) than by two other candidate proxy measures derived from structural MRI. **(a)** Correlation between hierarchy and cortical thickness. **(b)** Correlation between hierarchy and geodesic distance from primary visual cortex (V1), which is a rostro-caudal gradient. **(c)** Comparison of hierarchy correlation values for the myelin map, thickness map, and distance from V1. The myelin map is much more strongly correlated with hierarchy than the other two maps (*P <* 10^*−*3^). Statistical significance is calculated by a test of the difference between dependent correlations (*, *P <* 10^*−*1^; **, *P <* 10^*−*2^; ***, *P <* 10^*−*3^).

How well does the myelin map capture hierarchy relative to other putative proxy measures? We compared the performance of the myelin map against two alternative proxy candidates derived from structural MRI^19^: the map of cortical thickness, as cortex is generally thicker in association cortex than sensory cortex; and the map of geodesic distance from primary visual cortex, which defines a posterior-anterior gradient. We found that the myelin map was more predictive of hierarchy than were either of the two other candidate proxy measures (Fig. 2). The strong inverse relationship supports the cortical myelin map as a noninvasive proxy measure for hierarchy. The myelin map can be readily applied as a proxy measure for hierarchy in human cortex, for which lack of tract-tracing data precludes the direct characterization of hierarchy according to the conventional approach.

### Hierarchical gradients in cortical microcircuit specialization

To test for hierarchical specialization of microcircuit properties across human cortex, we examined areal patterns of cortical gene expression variation from the AHBA in relation to the myelin map. The AHBA is a transcriptional atlas that contains expression levels measured with DNA microarray probes and sampled from hundreds of neuroanatomical structures in the left cortical hemisphere across six normal post-mortem human brains^6^. From these data, we calculated group-averaged gene expression maps with 180 unilateral cortical areas using a multimodal parcellation from the Human Connectome Project^20^ (Fig. 3, see Methods). Due to the strong inverse relationship observed between the myelin map and hierarchy, if gene expression level is negatively correlated with myelin map value across areas, then expression level increases with position along the anatomical hierarchy (i.e., increasing from sensory to association cortex); conversely, a positive correlation indicates decreasing expression level along the hierarchy. To support the validity of our interpretations, we compared the myelin map correlation (MMC) of microcircuitry-related genes in human cortex to more direct anatomical measures in monkey cortex, with focus on cytoarchitecture, inhibitory interneuron densities, and synaptic processes (Fig. 4).

**Figure 3:**
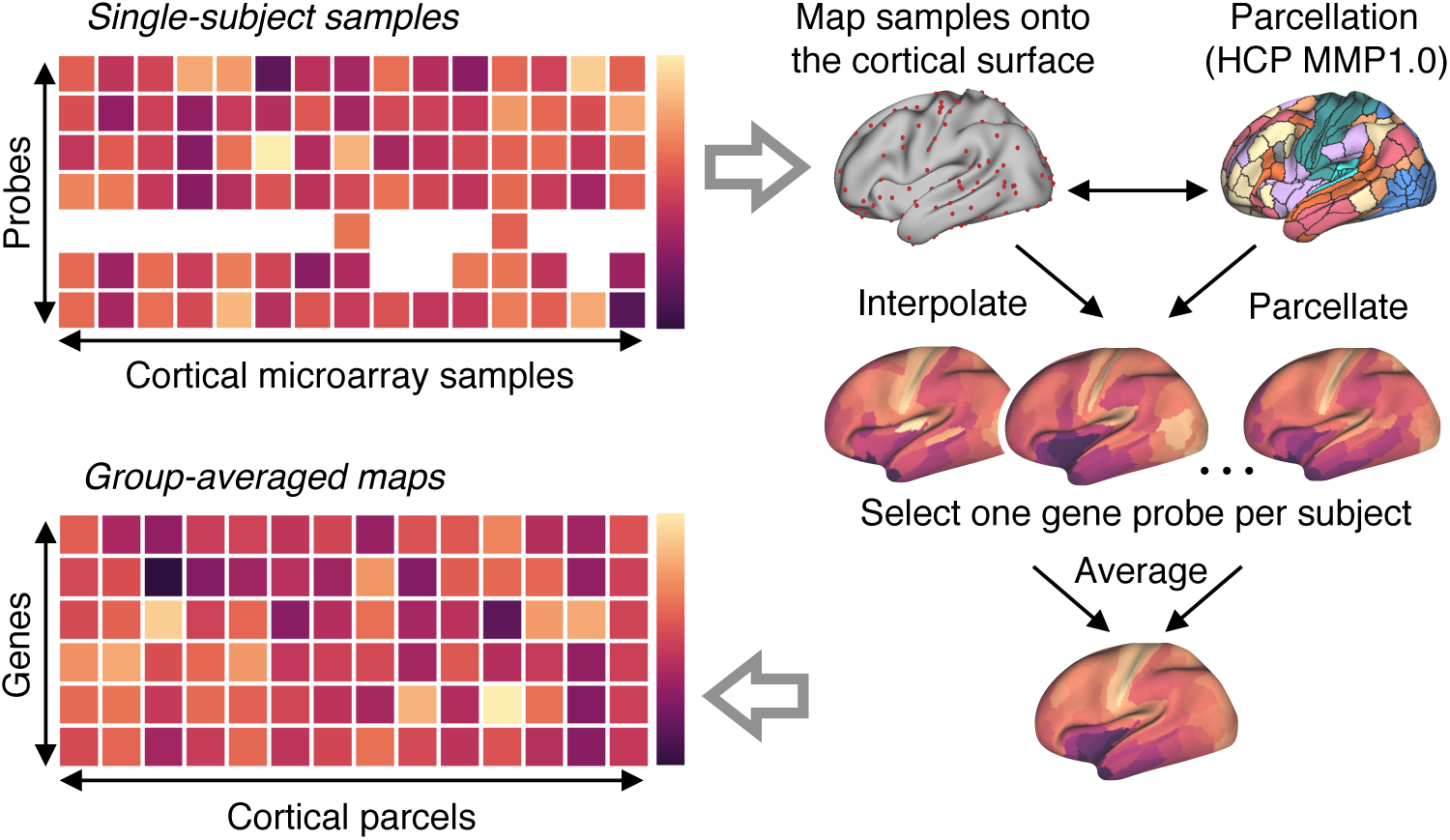
Procedure for generating group-averaged parcellated maps of gene expression levels. All analyses of gene expression patterns used group-averaged parcellated expression maps derived from the Allen Human Brain Atlas (AHBA) (see Methods for details). The AHBA contains genes expression levels measured with DNA microarray probes and sampled from hundreds of neuroanatomical structures in the left hemisphere across six normal post-mortem human brains. First, cortical samples were mapped from volumetric space onto a two-dimensional cortical surface, for each subject. Second, parcellated gene expression maps were constructed, for each subject, using a parcellation of the cortical surface into contiguous areas. For genes profiled by multiple microarray probes, we selected a single representative probe for each subject. Finally, a group-level parcellated expression map for each unique gene was computed by averaging parcellated expression levels across subjects’ selected gene probes.

**Figure 4:**
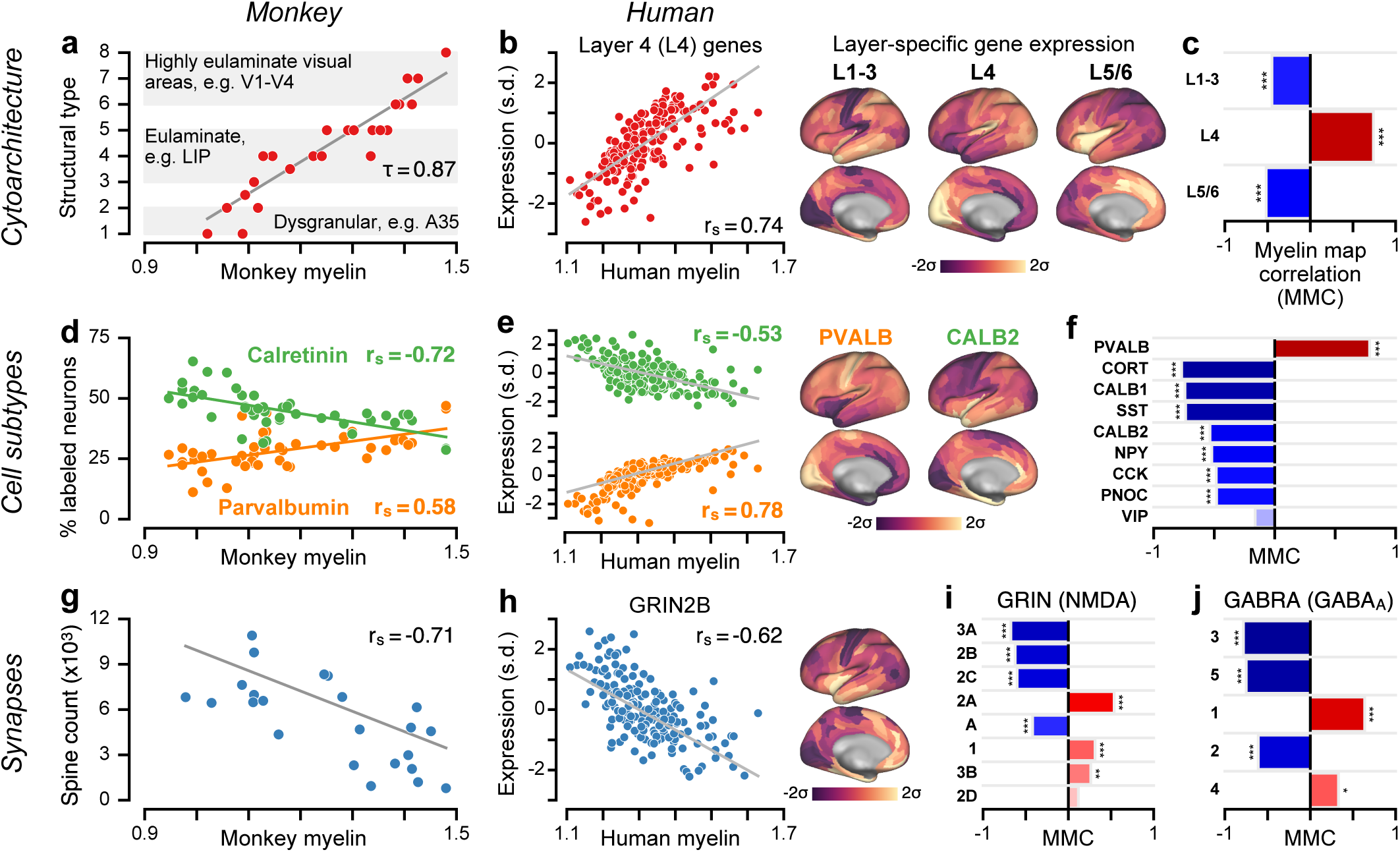
Myelin map topography capture specialization of cortical microcircuitry in humans and nonhuman primates. **(a)** Cytoarchitectural type is reliably predicted by the macaque monkey cortical myelin map (*τ* = 0.87, *P <* 10^*−*5^; Kendall’s tau correlation). **(b)** The average expression map of genes preferentially expressed in human granular layer 4 (L4) is positively correlated with the human cortical myelin map (*τ*_*s*_ = 0.74, *P <* 10^*−*5^; Spearman rank correlation), consistent with a more prominent granular L4 in sensory than association cortex. **(c)** Average expression maps of laminar-specific genes show significant myelin map correlations (MMCs). L1-3: supragranular layers 1-3; L5/6: infragranular layers 5 and 6. **(d)** The monkey cortical myelin map captures areal variation in the relative proportions of calretinin-and parvalbumin-positive inhibitory interneurons. **(e)** Genes coding for calretinin (*CALB2*) and parvalbumin (*PVALB*) exhibit homologous hierarchical gradients in human cortex. **(f)** MMCs of genes coding for markers of specific inhibitory interneuron cell types. **(g)** Basal-dendritic spine counts on pyramidal cells are significantly anti-correlated with the monkey myelin map (*τ*_*s*_ = −0.71, *P <* 10^*−*4^). **(h)** The gene coding for the NMDA receptor subunit NR2B (*GRIN2B*) exhibits a negative MMC (*τ*_*s*_ = −0.62, *P <* 10^*−*4^). **(i, j)** MMCs of genes coding for distinct subunits of the excitatory NMDA receptor and inhibitory GABA_A_ receptor. Statistical significance is calculated through a spatial autoregressive model to account for spatial autocorrelation (*, *P <* 10^*−*1^; **, *P <* 10^*−*2^; ***, *P <* 10^*−*3^).

An established feature of microcircuit specialization that varies along cortical hierarchy is the degree of laminar differentiation in local cytoarchitecture^21^: primary sensory cortex is highly laminated and exhibits a thick and well-defined granular layer, whereas association cortex is characterized by decreasing laminar differentiation and a gradual loss of the granular layer with progression along the hierarchy. In monkey cortex, we found a very strong correlation between myelin map value and cytoarchitectural type^21^ (Fig. 4a). In human cortex, we examined average expression profiles of genes reported to be preferentially expressed in specific cortical layers^22^. Consistent with trends observed in monkey cortex, we found a positive MMC for granular (L4) layer-specific genes, and negative MMCs for supra-(L1–3) and infra-granular (L5/6) layer-specific genes (Fig. 4b,c). These findings demonstrate that the noninvasive myelin map captures anatomical gradients related to cortical hierarchy in humans and nonhuman primates.

To gain further insight into microcircuit bases of hierarchical specialization, we examined the spatial distributions of markers for different inhibitory interneuron cell types. Inhibitory interneuron cell types fall into several biophysically distinct classes which differ in their synaptic connectivity patterns, morphology, electrophysiology, and functional roles^23,24^. In monkey cortex, we found that immunohistochemically measured densities of parvalbumin- and calretinin-expressing interneurons exhibit positive and negative MMCs, respectively (Fig. 4d). Consistent with these results, in human cortex we found corresponding hierarchical gradients in the expression profiles for the genes which code for parvalbumin and calretinin (Fig. 4e). In general, we observed strong hierarchical gradients in transcriptional markers for a number of inhibitory interneuron cell types^23^ (Fig. 4f), as well as for composite gene expression profiles associated with specific neuronal cell types derived from RNA sequencing in individual human neurons^25^ (Extended Data Fig. 3). These findings suggest that hierarchical gradients in neuronal cell-type distributions may contribute to specialization of cortical microcircuit function.

Gradients in the composition of synapses may endow cortical areas with diverse physiological properties required to perform the various computations which underlie specialized cognitive and behavioral functions. One putative microanatomical correlate for the strength of recurrent synaptic excitation in local cortical microcircuits is the number of excitatory synapses on pyramidal neurons, which can be quantified by counting the number of spines on pyramidal cell dendrites. In monkey cortex, we found a strong negative MMC for basal-dendritic spine counts on cortical pyramidal neurons^26^ (Fig. 4g). This finding suggests a gradient of increasing local recurrent excitation strength along the cortical hierarchy in primates^18^.

Distinct subunits of synaptic receptor proteins that mediate neurotransmission are differentially expressed across neuronal cell types and produce physiologically diverse synaptic properties. In the AHBA dataset, we examined expression profiles of genes that code for various excitatory and inhibitory synaptic receptor subunits (Fig. 4h–j). The gene *GRIN2B*, which codes for a glutamatergic NMDA receptor subunit mediating local synaptic excitation preferentially in primate association cortex^27^, exhibited a strong negative MMC, suggesting increased recurrent excitation strength in association cortical areas and consistent with the spine count gradient observed in monkey. Gene sets coding for neuromodulatory synaptic receptor subunits also contains strong positive and negative hierarchical gradients (Extended Data Fig. 4). The positive and negative MMCs reported in Fig. 4i,j suggest that gradients in local excitatory and inhibitory synaptic machinery contribute to the functional specialization of cortical microcircuitry^4,18^.

### Hierarchy captures the dominant axis of transcriptomic variation

How well does the myelin map capture areal variation in the transcriptomic architecture of human cortex in general? We performed principal component analysis (PCA) to identify the dominant areal patterns underlying gene expression variation (Fig. 5a–e, Extended Data Fig. 6). To test for generality of effects, we analyzed categorical sets of genes which are preferentially expressed in human brain tissue, neurons, oligodendrocytes, and synaptic compartments^28,29^. To assess statistical significance of effects, we developed a method for surrogate testing using randomized maps that preserve the spatial autocorrelation structure of the myelin map (Extended Data Fig. 7, see Methods). The first principal component (PC1) is the spatial map that captures the maximal amount of overall variance in gene expression across areas (Fig. 5a). Across all five gene sets, PC1 captures a large fraction of gene expression variance (range: 22–28%, more than twice PC2) (Fig. 5b, Extended Data Fig. 6), revealing that cortical gene expression patterns are effectively low-dimensional. Moreover, PC2 and PC3 consistently fractionated early sensory cortical areas by sensory modality, separating somatomotor cortex from early visual cortex across all five gene sets (Extended Data Fig. 5).

**Figure 5:**
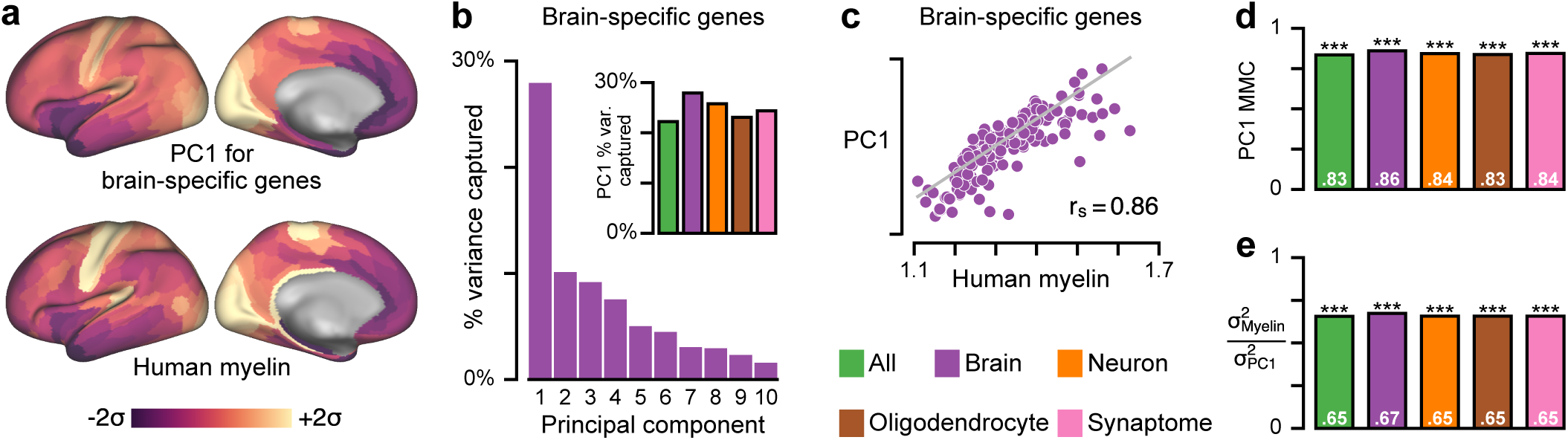
Myelin map topography captures the dominant axis of gene expression variation across human cortex. **(a)** The first principal component (PC1), here for a set of brain-specific genes, is the areal map that linearly captures the maximum variation in gene expression. **(b)** PC1 captures a large fraction of total gene expression variance. *Inset:* Variance captured by PC1 for five gene sets: all genes, and genes preferentially expressed in brain, neurons, oligodendrocytes, and synaptic processes. **(c)** PC1 for this gene set is highly correlated with the myelin map (MMC = 0.86; *P <* 10^*−*4^). **(d)** Across all sets, PC1 exhibits a highly similar areal topography to the myelin map (MMC range: 0.84–0.86; *P <* 10^*−*4^ for each). **(e)** Gene expression variance captured by the myelin map 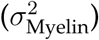 relative to PC1 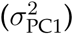. Statistical significance is calculated through permutation testing with surrogate maps that preserve spatial autocorrelation structure (*, *P <* 10^*−*1^; **, *P <* 10^*−*2^; ***, *P <* 10^*−*3^).

**Figure 6:**
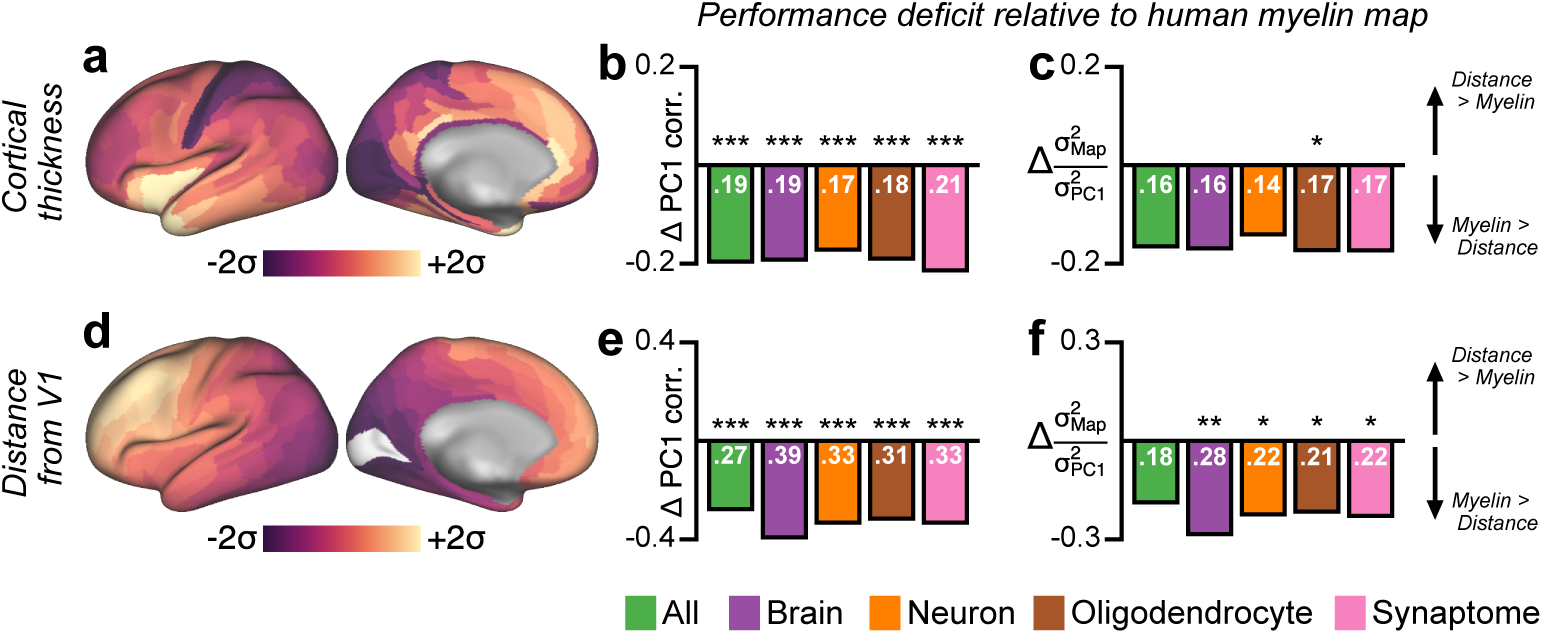
Principal component analysis (PCA) shows that the dominant mode of gene expression (PC1) is better captured by the myelin map than by other candidate proxies. **(a)** Parcellated map of human cortical thickness. **(b)** The difference in correlation with PC1 between the myelin map and the cortical thickness map, i.e., (*τ_s_*(Myelin, PC1) −*τ_s_*(Thickness, PC1)), across several categorical gene sets. Positive values indicate that the myelin map is more strongly correlated with PC1 than is the thickness map. Statistical significance is calculated by a test of the difference between dependent correlations (*, *P <* 10^*−*1^; **, *P <* 10^*−*2^; ***, *P <* 10^*−*3^). **(c)** The difference in the fraction of gene expression variance captured, relative to the variance captured by PC1, between the myelin map and the cortical thickness map, i.e., 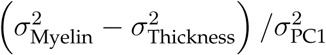, across several categorical gene sets. Positive values indicate that the myelin map captures more gene expression variance than does the thickness map. Statistical significance is calculated through permutation testing with surrogate maps that preserve spatial autocorrelation structure (*, *P <* 10^*−*1^; **, *P <* 10^*−*2^; ***, *P <* 10^*−*3^). **(d)** Parcellated map of geodesic distance from primary visual cortical area V1. **(e)** The difference in correlation with PC1 between the myelin map and the map of distance from area V1. **(f)** The difference in the fraction of gene expression variance captured, relative to the variance captured by PC1, between the myelin map and the map of distance from V1.

**Figure 7:**
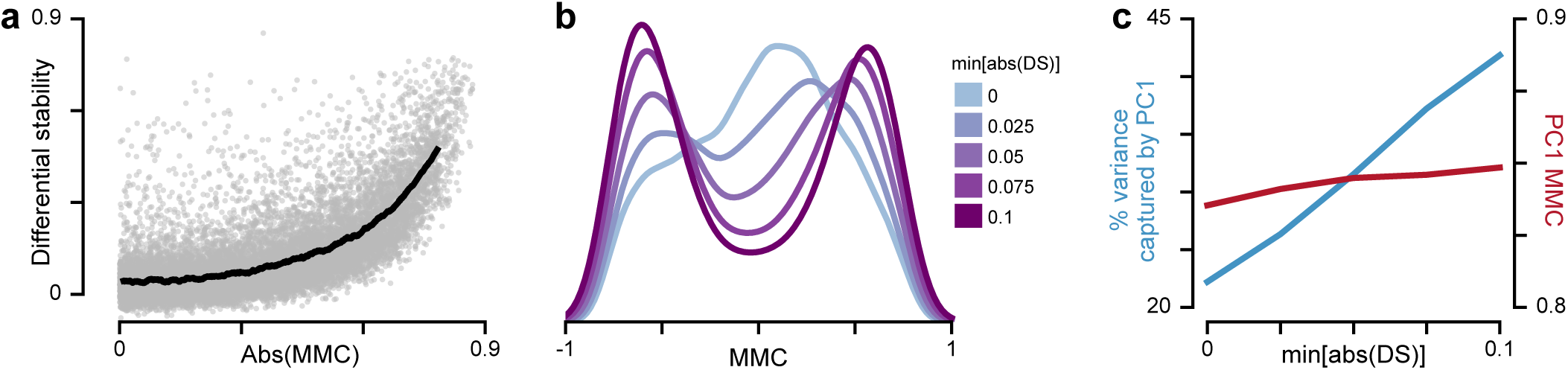
Expression profiles of genes which exhibit strong hierarchical gradients tend to be relatively stable across individuals. **(a)** Differential stability (DS), defined as the mean pairwise Spearman rank correlation between subjects’ expression maps, as a function of the magnitude of the myelin map correlation (MMC) (*τ*_*s*_ = 0.68, *P* < 10^*−*5^; Spearman rank correlation). Each gray dot represents a single gene. The black line indicates the average value in a sliding window of size 600 points. **(b)** Filtering genes by a threshold on DS alters the shape of the MMC distribution. Increasing the DS threshold filters out genes whose expression profiles are not relatively consistent across subjects. The trough which develops near MMC=0 suggests that high-DS genes preferentially exhibit strong hierarchical gradients. **(c)** Thresholding genes by DS substantially increases variance captured by the first principal component (PC1) of gene expression variation (*blue*) and only marginally increases PC1’s MMC (*red*).

Remarkably, we found that myelin map topography is strongly correlated with PC1, i.e., the dominant axis of gene expression variation, across all gene sets (MMC range: 0.84–0.86; *P <* 10^*−*4^) (Fig. 5c,d). We can also quantify how much gene expression variance is captured by the myelin map (see Methods). We found that across all gene sets the myelin map captures roughly two-thirds as much variance as PC1, which by construction is the spatial map that captures the maximum possible gene expression variation (Fig. 5e). We compared performance of the myelin map against the two alternative candidate proxy maps, cortical thickness and geodesic distance from primary visual cortex (Fig. 6). Across all gene sets, the myelin map was more strongly correlated with PC1 and captured more gene expression variance than either alternative map. The close alignment between myelin map topography and gene expression variance suggests that the dominant axis of transcriptomic organization in human cortex relates to hierarchy.

Genes that are especially vital to normal healthy cortical function may be more likely to have consistent spatial expression profiles across individual subjects. Hawrylycz and colleagues defined differential stability (DS) as the mean pairwise correlation between subjects’ individual gene expression profiles, which they found predicts association with key neurobiological functions^7^. We found a strong nonlinear and positive relationship between cortical DS and MMC magnitude (Fig. 7a). To gain additional insight into this relationship, we explored how the MMC distribution is impacted by filtering genes through a DS threshold. Exclusion of low-DS genes alters the shape of the MMC distribution, from having a peak near zero to having a trough near zero and roughly symmetric bimodal peaks at strong MMCs (Fig. 7b). Furthermore, exclusion of low-DS genes renders gene expression patterns more quasi-one-dimensional, as it strongly increases the fraction of gene expression variance captured by PC1, while marginally increasing the similarity of PC1 with the myelin map (Fig. 7c). Together, these results suggest that high-DS genes, i.e., genes whose spatial expression maps are consistent across individuals, preferentially exhibit strong positive and negative hierarchical gradients.

### Hierarchically expressed genes are enriched for functional and disease annotations

To examine the functional roles of genes with strong hierarchical variation, we tested for their preferential enrichment in gene sets defined by functional and disease ontologies. We found that genes with stronger MMCs are enriched in more functional categories, relative to genes with weaker MMCs, for all functional gene ontologies tested^7,30^: biological processes, cellular components, molecular functions, microRNA binding sites, and drug targets (Fig. 8a). These results suggest that diverse key cell-biological processes contribute to hierarchical differentiation of cortical microcircuitry. Finally, we examined whether hierarchical expression is a preferential property of genes associated with psychiatric and neurological disorders. For instance, we found that the genes *APOE* and *SNCA*, which are strongly linked to Alzheimer’s and Parkinson’s diseases, respectively^31^, exhibit robust negative MMCs and are therefore more highly expressed in association cortex (Fig. 8b,c). For a systematic examination, we statistically quantified the enrichment of genes with strong hierarchical variation in disease-related gene sets^7^, obtained from the DisGeNet database^32^. We found that genes with strongly negative MMCs were significantly over-represented across multiple disease-related gene sets (Fig. 8d). In particular, gene sets for schizophrenia, bipolar disorder, autistic disorders, and depressive disorders are significantly enriched with strongly negative MMC genes which are more highly expressed in association cortex. These findings suggest that brain disorders involve differential impacts to areas along the cortical hierarchy.

**Figure 8:**
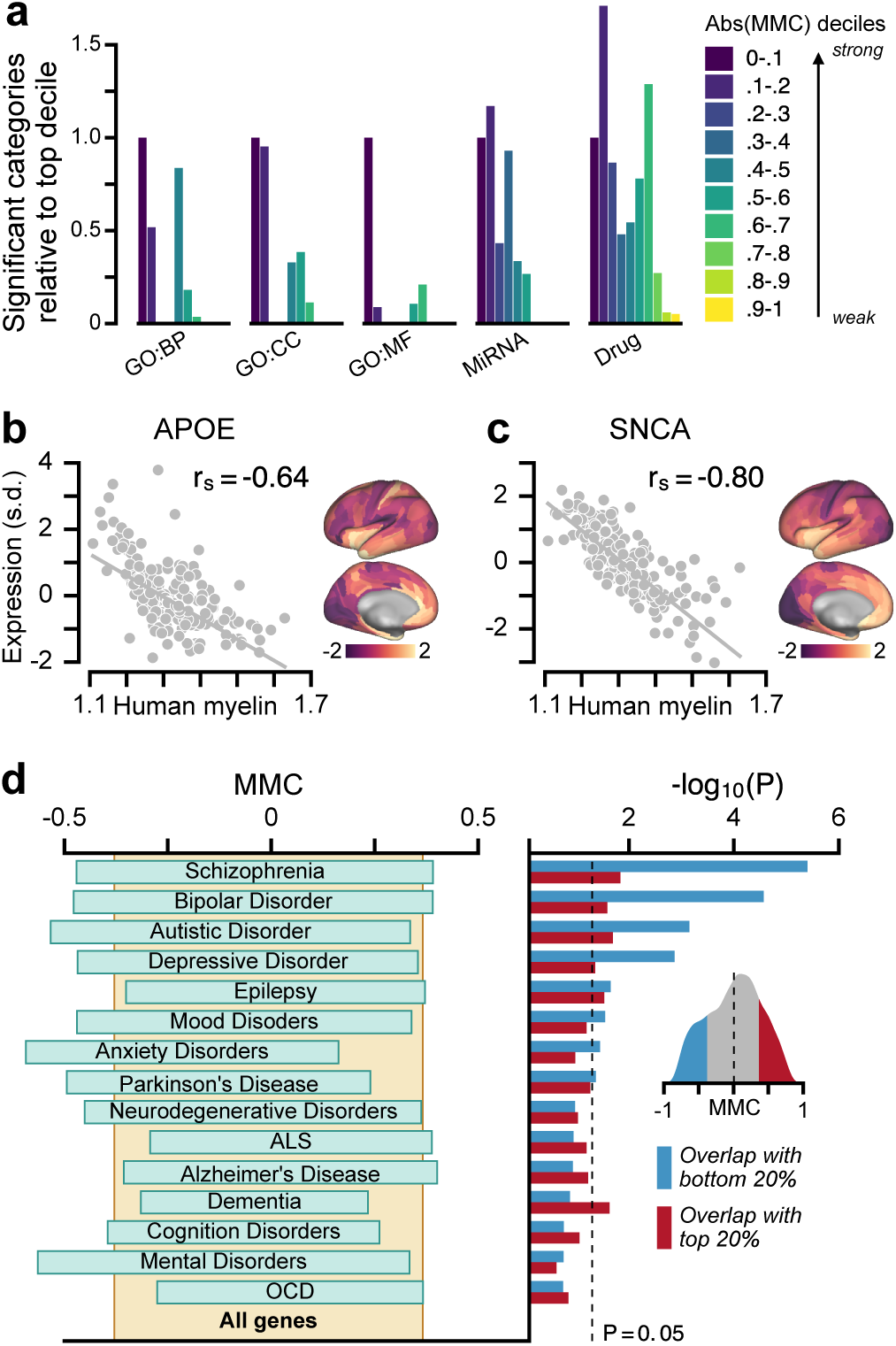
Hierarchical variation relates to enrichment in neurobiological function and brain disorders. **(a)** Genes with strong MMCs are overrepresented in functional annotations across multiple gene ontologies (GOs). BP, biological process; CC, cellular component; MF, molecular function; MiRNA, microRNA binding site. **(b, c)** Two key risk genes for neurodegenerative disorders, *APOE* for Alzheimer’s disease and *SNCA* for Parkinson’s disease, exhibit strongly negative MMCs, with higher expression levels in association cortex relative to sensory cortex (*APOE*: MMC = −0.64, *P <* 10^*−*15^; *SNCA*: MMC = −0.80, *P <* 10^*−*42^). *APOE* is a leading risk gene for Alzheimer’s disease. The *ε*4 allele of *APOE* is the largest genetic risk factor for late-onset Alzheimer’s disease. *SNCA* (*PARK1*/*PARK4*) is a key risk gene for Parkinson’s disease. Duplication of *SNCA* is risk factor for familial Parkinson’s disease with dominant inheritance. *SNCA* codes for the alpha-synuclein protein which is the primary component of Lewy bodies, a biomarker of Parkinson’s disease. **(d)** Genes with strong negative MMCs are overrepresented in multiple gene sets associated with neuropsychiatric disorders. *Left panel:* 20–80% ranges of MMC for gene sets. *Right panel:* Enrichment is quantified by the hypergeometric test, which assesses the statistical significance of overlap between each gene set and the top (red) or bottom (blue) 20% MMC genes. *Inset:* Distribution of MMCs across genes.

## Discussion

Taken together, our findings show that cortical hierarchy provides an organizing principle for the transcriptomic architecture of human cortex. First, the MRI-derived myelin map provides a noninvasive neuroimaging proxy for anatomical hierarchy in the absence of axonal tract-tracing data. Second, the principal axis of transcriptional variation across human cortex aligns with cortical hierarchy as captured by the myelin map. Third, this hierarchical axis reflects microcircuit specialization involving synapses and cell types, with relevance to brain disease pathophysiology. Strong similarities between the patterns of anatomical, functional, and transcriptional variation suggest that hierarchical gradients of microcircuit properties play key roles in the functional specialization of large-scale networks across the human cortex.

Specialization of cortical function may derive in part from the multiple features of microcircuitry identified here to exhibit hierarchical gradients. In addition to variation in synaptic subunit composition, stronger recurrent excitation in association cortex can endow association circuits with longer intrinsic timescales of spontaneous activity^18,33^, as observed empirically^4,5^, which subserve the prolonged integration of signals. Furthermore, computational modeling of cortical circuits identifies recurrent excitation strength as a key property governing functional specialization across areas for core cognitive computations such as working memory and decision making^33,34^. Hierarchical gradients of inhibitory interneuron cell types can additionally shape interareal specialization of dynamics and function, due to cell-type differences in physiology and synaptic connectivity^23,24^. For example, parvalbumin-expressing inhibitory interneurons preferentially target the perisomatic areas of pyramidal neurons where they can gate pyramidal-neuron outputs. In contrast, calretinin-expressing inhibitory interneurons preferentially target distal dendrites of pyramidal neurons and other inhibitory interneurons, where they may play key computational roles in disinhibition-mediated gating of dendritic inputs^35^.

Our study adds to a growing understanding of how transcriptomic specialization shapes cortical function. Transcriptional diversity, particularly of genes which regulate synaptic function and ion channel activity, relates to the spatiotemporal organization of intrinsic activity in large-scale cortical networks^7,9–11^, and transcriptional markers for synaptic, neuronal, and axonal structure relate to patterns of anatomical connectivity^11,12^. Of note, Hawrylycz et al. (2015) found that genes most strongly predictive of functional connectivity patterns in cortex were shifted toward high DS, and that high-DS genes were significantly enriched in gene sets related to functional ontologies and brain diseases, leading the authors to suggest these genes constitute a “canonical transcriptional blueprint” for the human brain^7^. We found that cortical DS is strongly associated with MMC (Fig. 7a), and that high-MMC genes exhibit similar functional and brain disease-related enrichments, indicating that genes whose expression topography is highly conserved across individuals preferentially exhibit strong hierarchical gradients in cortex.

Our findings show that the myelin map generally captures an axis of hierarchical differentiation across cortex that reflects multiple features of interareal variation beyond just intracortical myelin content. Intracortical myelination itself may contribute to hierarchical functional specialization in multiple ways^36^. First, higher myelination may support speed, fidelity, and efficiency of axonal transmission in sensory cortex^37^. Second, myelination may regulate plasticity through exposure of axons for synapse formation^38^, leading to greater plasticity in lightly-myelinated association cortex than in heavily-myelinated sensory cortex. Third, myelination may reflect proportions of cell types, such as heavily myelinated parvalbumin-expressing interneurons^39^. We note that there are interesting deviations between the topographies of the myelin map and other hierarchical features. For instance, primary motor cortex and retrosplenial cortex exhibit high myelin map values^17^ yet differ from primary sensory areas in their laminar structure.

Our findings have implications for computational models of large-scale dynamics in human cortex, which are applied to explain how the structure of resting-state functional connectivity emerges through long-range interactions among cortical areas. Leading circuit models of human cortex treat cortical microcircuitry as homogeneous across cortex, with all cortical areas modeled as nodes with identical properties^40^. Recent modeling has shown that hierarchical heterogeneity can shape functional connectivity^18^, and its alterations in disease states^41^. Empirical resting-state functional connectivity exhibits structure related to cortical hierarchy^42^, which is captured by myelin map topography^43^. We propose that interareal variation of microcircuit parameters can be anatomically constrained by structural neuroimaging maps, such as the myelin map, in next-generation circuit models of human cortex.

Multiple lines of evidence point to a transcriptional basis for disease phenotypic variation, linking white matter dysconnectivity^44^ and developmental changes in structural topology^45^ to genes implicated in schizophrenia. Further characterization of the developmental trajectory of hierarchical transcriptomic specialization^44,46,47^ may inform the progression of neurodevelopmental disorders. Hierarchical gradients in drug targets, such as receptor subunits, enables preferential modulation of sensory or association cortical areas through pharmacology, which may guide future rational design of treatments to target specific macroscale cortical circuits. Large-scale mapping of the cortical transcriptome at finer spatial resolution will further elucidate the microcircuit basis of hierarchical specialization with laminar^22^ and cell-type^8,25^ specificity.

## Acknowledgements

We thank B.D. Fulcher, X.-J. Wang, R. Chaudhuri, and D.C. Glahn for useful discussions. This research was supported by NIH grants R01MH112746, R01MH108590, and TL1TR000141, and BlackThorn Therapeutics.

## Footnotes

**Author Contributions** J.B.B., W.J.M., A.B., A.A. and J.D.M. designed the research. J.B.B., M.D., W.J.E., N.N., and L.J. analyzed the data. J.D.M. supervised the project. J.B.B. and J.D.M. wrote the manuscript and prepared the figures. All authors contributed to editing the manuscript. **Author Information** Correspondence and requests for materials should be addressed to J.D.M.

## Methods

### Parcellated cortical myelin maps (T1w/T2w)

Cortical myelin maps were defined as the ratio of T1- to T2-weighted (T1w/T2w) MRI maps as previously characterized^17,36^, using the surface-based CIFTI format^20^. The T1w/Tw2 map has been shown to correlate with grey-matter intracortical myelination and to reflect architectonic boundaries between cortical areas^17,36^. Of note, it may not index myelin content in white matter. The group-averaged (*N* = 69) human myelin map was obtained from the publicly available Conte69 dataset, which was reported previously to study myelin maps^17^. The group-averaged (*N* = 334) cortical thickness map was obtained from the Human Connectome Project (HCP)^48^. Human myelin map values for the left cortical hemisphere were parcellated into 180 areas using the Multi-Modal Parcellation (MMP1.0) from the HCP^20^. Assignment of MMP1.0 parcels to functional networks (Fig. 1b, Extended Data Fig. 1c) was performed through community detection analysis[49] on time-series correlation from the HCP resting-state fMRI dataset.

The group-averaged myelin (T1w/Tw) map and thickness map for macaque monkey cortex were obtained from the publicly available BALSA database^50^ (*N* = 19) (https://balsa.wustl.edu/study/show/W336). Monkey myelin map values for the left cortical hemisphere were parcellated into 91 areas using the M132 parcellation which was used for the anatomical tract-tracing dataset^50^. Geodesic distance between two parcels *i* and *j* is calculated as the average of all pairwise surface-based distances between grayordinate vertices in parcel *i* and vertices in parcel *j*.

### Anatomical hierarchy levels in monkey cortex

To assess whether macaque cortical myelin maps could reliably capture the laminar-specific interareal projection patterns conventionally used to define anatomical hierarchy, we fit a generalized linear model (GLM) to quantitative laminar projection data, yielding ordinal hierarchy values in 89 cortical areas, following the procedure of ref. [15]. Anatomical tract-tracing data, derived from retrograde tracers, was obtained from the publicly available Core-Nets database (http://core-nets.org). Retrograde tracer was injected into a target area *i*, and the number of labeled neurons in source area *j* were counted. The fraction of external labeled neurons, *FLNe*_*ij*_, is a quantitative measure of connection strength defined as the number of labeled neurons in the source area normalized by the total number of labeled neurons in all external cortical source areas for a given injection^51^. Labeled neurons in the source areas are classified by location in either supragranular or infragranular layers. For a given projection, the proportion of supragranular labeled neurons, *SLN _ij_*, is defined as the ratio of *N*_supra_ to *N*_supra_ + *N*_infra_ for neurons labeled in source area *j*. As feedforward and feedback connections preferentially originate in supragranular and infra-granular layers, respectively^13–15^, SLN is a quantitative measure of hierarchical distance between two cortical areas^15^: under this paradigm for laminar-specific projection motifs, a pure feedforward connection from source area *j* to target area *i* would originate entirely in the superficial layers, resulting in an SLN of 1. Conversely, a pure feedback projection originating entirely in deep infragranular layers would result in an SLN of 0.

The GLM procedure for fitting hierarchy from SLN data is described in detail in ref. [15]. In brief, the hypothesis that SLN is indicative of hierarchical distance can be expressed as *g*(*SLN*_*ij*_) = *H_i_ − H_j_*, where *H*_*i*_ corresponds to the hierarchical position of area *i*, and *g* is an arbitrary and possibly nonlinear function linking SLN values on the unit interval (0, 1) to their corresponding hierarchical distance. We used a logit link function to map SLN values from the unit interval to the entire real number line following the procedure of ref. [18]. Fitting linear predictors (i.e. hierarchical levels) to logit-transformed SLN values formulates a type of generalized linear model, with maximum likelihood estimation assuming a binomial family probability distribution for the supra- and infragranular neuron counts. To assign more weight to stronger connections during model estimation of hierarchical levels, we also weight each pathway in the model by the negative logarithm of the FLNe value. We clip SLN values to lie in the interval (0.01, 0.99) so the logit-transformed SLN value is well-defined for all pathways used to fit the model. Furthermore, to reduce the impact of noise on model parameter estimation, we only included pathways which contained at least 100 projection neurons when fitting the GLM; we confirmed that results were generally robust to the choice of neuron count threshold.

Maximum likelihood estimation of model parameters was done in the R programming language using the glm function. The model-estimated hierarchy levels, invariant under linear transformations, were rescaled to span the unit interval [0, 1]. To assess the statistical relationship between myelin map value and hierarchy level, we calculated the Spearman rank correlation between the 89 ordinal hierarchy values and their corresponding parcellated myelin map values (Fig. 1f). For visual clarity in Fig. 1c,d we remove this nonlinear transformation by displaying model-estimated hierarchy levels after applying the inverse-logit (i.e., logistic) transformation. This rescaling preserves the ordering of areas and therefore does not affect the reported Spearman rank correlations.

### Macaque monkey anatomical data: cytoarchitectural types, inhibitory interneuron densities, and pyramidal neuron spine counts

To quantify the statistical relationship between myelin map value and categorical cytoarchitectural type (Fig. 4a), we compared myelin map values to structural classification values reported for 29 regions of primate visual cortex, obtained from ref. [21]. To characterize hierarchical distributions of cortical inhibitory interneuron cell types (Fig. 4b), we compiled, from multiple immunohistochemical studies, the relative densities of inhibitory interneurons which are immunoreactive (ir) to the three calcium-binding proteins parvalbumin (PV), calretinin (CR), and calbindin (CB)^52–55^. To characterize hierarchical variation in pyramidal neuron excitatory synaptic connectivity (Fig. 4c), we compiled, from multiple studies by Elston and colleagues^56–61^, the number of spines of basal-dendritic trees of layer-3 pyramidal neurons.

For each of these three analyses, we produced a mapping between the 91 areas in the M132 atlas parcellation, where the myelin map values are calculated, to the architectonic areas reported in these collated studies (Supplementary Table 1). Where the anatomical mapping was not a one-to-one correspondence, we mapped the reported anatomical area onto the set of all M132 parcels with nonzero spatial overlap, and the myelin map value was calculated as the average across these M312 parcels.

### Gene expression preprocessing

The Allen Human Brain Atlas (AHBA) is a publicly available transcriptional atlas containing gene expression data, measured with DNA microarrays, that are sampled from hundreds of histologically validated neuroanatomical structures across six normal post-mortem human brains^6^. After no significant interhemispheric transcriptional differences were observed in the first two bilaterally profiled brains^6^, the remaining four donor brains were profiled only in the left cortical hemisphere^7^. To construct parcellated group-averaged expression maps, we therefore restricted all analyses to microarray data sampled from the left cortical hemisphere in each of the six brains. Microarray expression data and all accompanying metadata were downloaded from the AHBA (http://human.brain-map.org)^6,7^. The raw microarray expression data for each of the six donors includes expression levels of 20,737 genes, profiled by 58,692 microarray probes. These data were preprocessed according to following procedure:

1. Gene probes without a valid Entrez Gene ID were excluded.
2. Cortical samples exhibiting exceptionally low inter-areal similarity were excluded. We first computed the spatial correlation matrix of expression values between samples using the remaining 48,170 probes, then summed this matrix across all samples. Samples whose similarity measure was more than five standard deviations below the mean across all samples were excluded. At most, this step excluded three samples within a subject.
3. Samples whose annotations did not indicate that they originated in the left hemisphere of the cerebral cortex were excluded. To focus analysis to neocortex, we also excluded samples taken from cortical structures that are cytoarchitecturally similar to the hippocampus, including the rhinal sulcus, piriform cortex, parahippocampal gyrus, and the hippocampal formation.
4. The remaining cortical samples were mapped from volumetric space onto a two dimensional cortical surface by minimizing the pairwise 3D Euclidean distance between the stereotaxic MNI coordinates reported for each sample, and each grayordinate vertex in the group-averaged surface mesh of the midthickness map in the Conte69 brain atlas. Cortical samples whose Euclidean distance to the nearest surface vertex was more than two standard deviations above the mean distance computed across all samples were excluded (excluding between 4 and 13 samples per subject). An average of 203 *±* 32 samples per subject, yielding 1219 total samples across all six subjects, remained at this stage.
5. Expression profiles for samples mapped onto the same surface vertex were averaged. Then expression profiles for each remaining sample were z-scored across gene probes.
6. Expression profiles for each of the 180 unilateral parcels in the HCP’s MMP1.0 cortical parcellation^20^ were computed in one of the two following ways. (I) For parcels which had at least one sample mapped directly onto one of their constituent surface vertices, parcellated expression values were computed by averaging expression levels across all samples mapped onto the parcel. (II) For parcels which had no samples mapped onto any of their constituent vertices, we first created densely interpolated expression maps, in which each surface vertex was assigned the expression level associated with the most proximal surface vertex onto which a sample had been mapped (i.e., a Voronoi diagram), determined using surface-based geodesic distance along the cortical surface; the average of expression levels across parcels’ constituent vertices was then computed to obtain parcellated expression values.
7. A coverage score was also assigned to each gene probe, defined as the fraction of 180 parcels that had at least one sample mapped directly onto one of its constituent surface vertices. Probes with coverage below 0.4 (i.e., probes for which fewer than 72 of the 180 parcels contained samples) were excluded.
8. For each gene profiled by multiple gene probes, we selected and used the expression profile of a single representative probe. If two probes were available, we selected the probe with maximum gene expression variance across sampled cortical structures, in order to more reliably capture spatial patterns of areal heterogeneity. If three or more probes were available, we selected a probe using a procedure similar to the one described in step 2: we computed a correlation matrix of parcellated gene expression values across the available gene probes, summed the resultant matrix along one of its dimensions to obtain a quantitative similarity measure for each probe, relative to the other gene probes, and selected the probe with the highest similarity measure, as it is most highly representative among all available gene probes.
9. Each subject-level gene expression profile was z-scored before we computed group-level expression profiles, which were obtained by computing the mean across subjects which were assigned a probe for that gene. Genes were excluded if fewer than four subjects were assigned a probe. Finally, group-level expression profiles were z-scored across areas for each gene.

These steps yielded group-averaged expression values for 16,040 genes across 180 cortical areas, which were used for all analyses reported here. The myelin map correlation (MMC) for each gene is reported in Supplementary Table 2.

### Categorical gene sets

We conducted analyses on biologically and physiologically meaningful gene sets extracted from existing databases and neuroscientific literature, reported below (Supplementary Table 2):

1. **Brain-specific.** Genes with expression specific to human brain tissue, relative to other tissues, were obtained from supplementary data set 1 of ref. [62]. Following ref. [28], brain-specific genes were selected for which expression in brain tissue was four times higher than the median expression across all 27 different tissues.
2. **Neuron- and oligodendrocyte-specific.** Brain genes with expression specific to neurons or oligodendrocytes, relative to other central nervous system (CNS) cell types, were obtained from supplementary data set S3b of ref. [63]. Following ref. [28], neuron-specific genes were selected for which log-expression in neurons of P7n cell type in the mouse was 0.5 greater than the median log-expression across 11 CNS cell types.
3. **Synaptome.** Four synaptic gene sets encoding proteins in the presynaptic nerve terminal, presynaptic active zone, synaptic vesicles, and postsynaptic density, were obtained from SynaptomeDB, an ontology-based database of genes in the human synaptome^29^.
4. **Neuron subtype-specific.** Gene sets representing distinct classes of neuronal subtypes were obtained from ref. [25], in which clustering and classification analyses yielded 16 distinct neuron subtypes, on the basis of differential gene expression measured by RNA sequencing from single neurons in human cortex. The fraction of positive values using exon-only derived transcripts per million (TPM) associated with each subtype-specific gene were obtained from supplementary table S5; within each neuronal subtype cluster, the TPM values for the cluster genes were normalized and used to create a weighted gene expression profile representative of each subtype’s spatial topography (Extended Data Fig. 3).
5. **Layer-specific.** Sets of laminar-specific genes localized to different layers of human neocortex were obtained from supplementary table S2 of ref. [22]. Genes were broadly grouped into sets representative of supragranular (L1–3), granular (L4), and infragranular (L5/6) layers.

### Spatial autoregressive modeling

Significance values as indicated by the number of stars reported on barplots for myelin map correlations were corrected to account for spatial autocorrelation structure in parcellated myelin map and gene expression values. Because physical quantities like cortical myelination and gene expression must vary smoothly and continuously in space, measurements recorded from proximal cortical areas tend to be more similar than measurements recorded from distal areas of cortex. This departure from the assumption of independent observations biases calculations of statistical significance. To model this spatial autocorrelation, we used a spatial lag model (SLM) commonly applied in the spatial econometrics literature^64^, of the form *y* = *ρW y* + *Xβ* + *ν*, where *W* is a weight matrix implicitly specifying the form of spatial structure in the data, and *ν* is normally distributed.

To implement a spatial lag model in the python programming language, we used the maximum likelihood estimation routine defined in the Python Spatial Analysis Library (*pysal*)^65^. We first determined the surface-based spatial separation between each pair of cortical parcels by computing the mean of the pairwise distances between a vertex in parcel *i* and a vertex in parcel *j*, from which we constructed a pairwise parcel distance matrix, *D*.

Similarity of gene expression profiles was well-approximated by an exponential decaying spatial autocorrelation function (Extended Data Fig. 7), as was found in mouse cortex^1-2^. We fitted the correlation of gene expression profiles between two areas with the exponential function Corr(*x_i_, x_j_*) *∼* exp(*−D_ij_/d*_0_), where *x*_*i*_ and *x*_*j*_ are vectors containing the parcellated gene expression values at parcels *i* and *j*, *D*_*ij*_ is the geodesic distance between the parcels, and *d*_0_ is the characteristic spatial scale of autocorrelation. We empirically determined *d*_0_ by first constructing the pairwise gene co-expression matrix *C*_*ij*_ = Corr(*x_i_, x_x_*), where *x*_*i*_ and *x*_*j*_ are vectors containing the parcellated gene expression values at parcels *i* and *j*. We then fit the free parameter *d*_0_ using ordinary least squares (OLS) regression on the off-diagonal (upper-triangular) elements of the gene co-expression and parcel distance matrices, so as to minimize the sum-of-squared-residuals between empirical and model-estimated gene co-expression values over all pairs of cortical parcels, 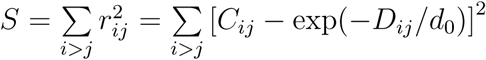. This empirical fit was performed on a set of brain-specific genes. Using the OLS estimate of the spatial autocorrelation scale from the fit to the empirical gene expression data, we calculated the elements of the spatial weight matrix, *W*_*ij*_ = exp(*−D_ij_/d*_0_). Finally, we fit the SLM to parcellated gene expression profiles, using the maximum likelihood estimator routine (pysal.spreg.ml_lag.ML_Lag) in *pysal*. P-values indicated by the number of stars in the bar plots of myelin map correlation correspond to p-values for model parameter *β* defined above.

Of note, spatial autoregressive model parameters do not have the same interpretation as they do in OLS regression. The parameter *β* reflects the direct (i.e. local) impact on the dependent variable *y* due to a unit change in the independent variable *x*. In addition, because of the underlying spatial structure, the direct impact of *x*_*i*_ on *y*_*i*_ results in an indirect effect of *y*_*i*_ on neighboring *y*_*j*_. Therefore *β* cannot be interpreted as a corrected, global correlation coefficient, and we restrict our use of the SLM to correcting for the biasing effect of spatially autocorrelated samples on reported significance values.

### Theil-Sen estimator

Trend lines in figures are calculated by the Theil-Sen estimator, which is a nonparametric estimator of linear slope, based on Kendall’s tau rank correlation, that is insensitive to the underlying distribution and robust to statistical outliers^66^. It is defined as the median of the set of slopes computed between all pairs of points.

### Principal components analysis

We used principal component analysis (PCA) to identify the dominant modes of spatial variation in the transcriptional profiles of gene expression in the human cortex. For a set of *N* genes, each with group-averaged expression values for *P* cortical parcels, we constructed a gene expression matrix **G** with one row for each cortical parcel and one column for each unique gene (i.e. with dimensions *P × N*). The *P × P* spatial covariance matrix **C** was constructed by computing the covariance between vectors of gene expression values for each pair of cortical parcels: *C*_*ij*_ = Cov(*G_i_, G_j_*), where *G*_*i*_ is the *i*-th row in the matrix **G**, corresponding to the vector of *N* gene expression values for the *i*-th cortical parcel. Eigen-decomposition is performed on the spatial covariance matrix to obtain the matrix eigenvectors (i.e., the principal components, PCs) and their corresponding eigenvalues, which are the amount of variance captured by the corresponding PC. To enumerate each principal component, eigenvalues are ranked in descending order of absolute magnitude, with larger magnitudes indicating a greater proportion of the total variance captured by the associated PC (i.e., the associated mode of spatial covariation). PCA therefore allows for simultaneous identification of spatial patterns of covariation and quantification of the extent to which these spatial modes capture variance in cortical gene expression profile s.

To quantify the overlap of these spatial PCs with the cortical myelin map vector, we compute the Spearman rank correlation coefficient between each *P* -dimensional PC and the *P* -dimensional vector of myelin map values for each cortical parcel. We can quantify the amount of gene expression variance that is captured along any given spatial map, such as the myelin map (Fig. 5e, Extended Data Fig. 6g,k). From the spatial covariance matrix **C**, the variance captured along a unit-length vector **a**, here a demeaned and normalized map, is given by **a**^*⊤*^**Ca**.

### Surrogate data generation

To nonparametrically determine significance values in our PCA results, in Fig. 5 and Extended Data Fig. 6, we generated surrogate maps with a spatial autocorrelation structure matched to the empirical data (Extended Data Fig. 7b). Parameters characterizing the empirical spatial autocorrelation were determined numerically for the cortical myelin map, cortical thickness map, and the map of surface-based geodesic distance from area V1; in each case, we fit the data using a spatial lag model of the form **y** = *ρ***Wy**, where **y** is a vector of mean-subtracted map values. **W** is the weight matrix with zero diagonal and off-diagonal elements *W*_*ij*_ = exp(*−D_ij_/d*_0_), where *D*_*ij*_ is the surface-based geodesic distance between cortical areas *i* and *j*. Two free parameters *ρ* and *d*_0_ are estimated by minimizing the residual sum-of-squares^64^. Using best-fit parameter values 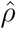 and 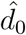, surrogate maps **y_surr_** are generated according to 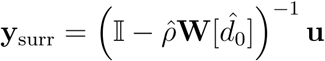, where **u** *∼ N* (0, 1). From these surrogate maps we construct null distributions for the appropriate statistics, and report significance values as the proportion of samples in the null distribution whose absolute value is equal to or greater than the absolute value of the test statistic.

### Functional enrichment analyses

Functional enrichments were determined using the ToppGene (https://toppgene.cchmc.org/) web portal^30^, including gene ontology annotations (biological process, cellular component, and molecular function); microRNA targets (from all sources indicated on https://toppgene.cchmc.org/navigation/database.jsp); and drug annotations (from DrugBank, Comparative Toxicogenomics Database, including marker and therapeutic, and Broad Institute CMAP). Significant genes in each category were identified using the ToppFun utility. Disease annotations were determined using curated disease gene associations in the DisGeNet database^32^ (http://www.disgenet.org/web/DisGeNET/menu/home). Hypergeometric testing was used to determine significant over-representation of brain-related disease genes in the top and bottom gene quintiles (20%, 3208 genes) ranked by myelin map correlation, following ref. [7].

---

**Extended Data Figure 1:**
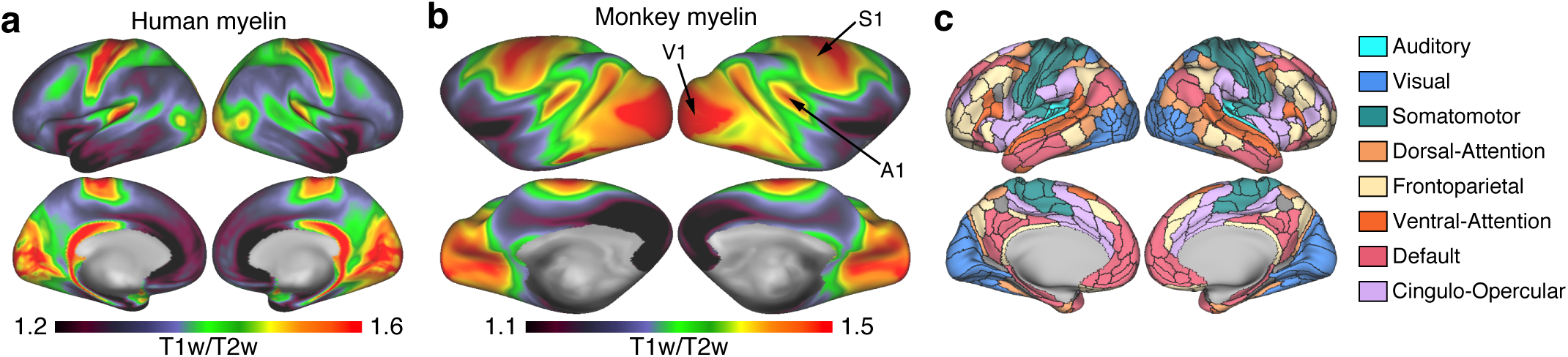
Cortical myelin maps exhibit inter-species homology and inter-hemispheric symmetry. **(a)** Unparcellated bilateral myelin map (T1w/T2w) in human cortex visualized on an inflated cortical surface. **(b)** Unparcellated bilateral myelin map (T1w/T2w) in monkey cortex visualized on an inflated cortical surface. Primary sensory areas (visual, V1; somatosensory, S1; auditory, A1) exhibit high myelin map values, as do their homologues in human cortex. **(c)** Functional networks derived from resting-state functional connectivity from the Human Connectome Project (HCP). Cortical areas are parcellated using the HCP multi-modal parcellation (MMP1.0). We assigned each region to a functional network using a community detection method applied to resting-state fMRI data from the HCP, and designated functional labels to networks, including three sensory and five association, that align with previously reported functional networks (with abbreviations labeled in Fig. 1b): Auditory (AUD), Visual (VIS), Somato- motor (SOM), Dorsal Attention (DAN), Frontoparietal (FPN), Ventral Attention (VAN), Default (DMN), and Cingulo-Opercular (CON).

**Extended Data Figure 2:**
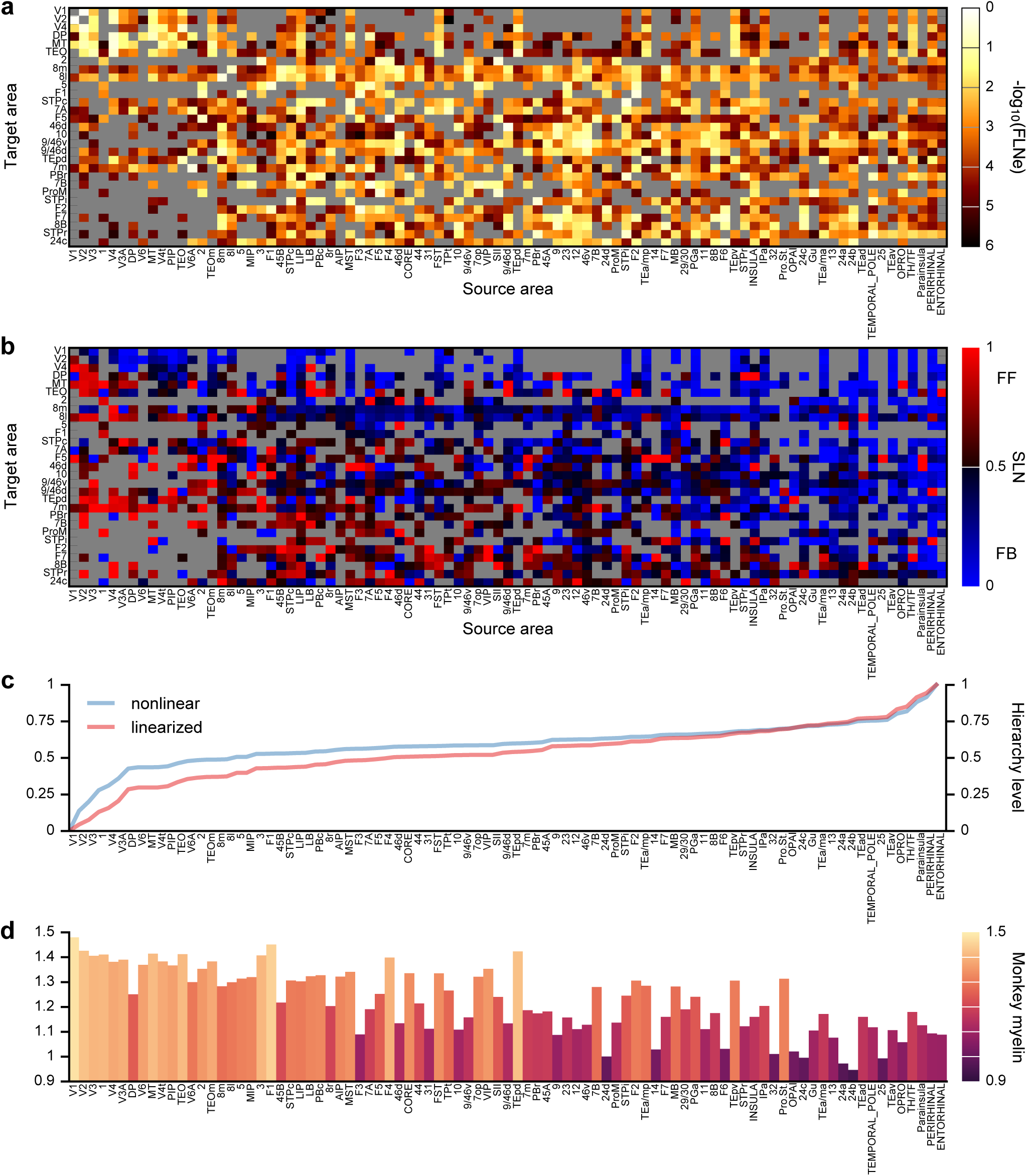
Anatomical cortical hierarchy derived from laminar-specific interareal projections in monkey cortex. **(a)** Fraction of external labeled neurons (*F LN e*). Target area *i* is injected with a retrograde tracer that labels neurons in many source areas; the *F LN e* in source area *j* is then defined as the fraction of all external labeled neurons terminating in area *i* that originated in source area *j*. Each row of the *F LN* matrix is therefore normalized to 1. Measurements which yielded no labeled neurons are marked in grey. **(b)** Fraction of supragranular layer neurons (*SLN*), defined as the fraction of neurons in an interareal projection (to target area *i* from source area *j*) originating in supragranular layers. An *SLN* of 1 indicates that all labeled projection neurons were of supragranular origin, reflecting a pure feedforward connection; an *SLN* of 0 indicates that all projection neurons originated in deep infragranular layers, reflecting a pure feed- back connection. Measurements which yielded no labeled neurons are marked in grey. **(c)** Model-estimated hierarchy values for 89 cortical regions. The blue line indicates hierarchy levels estimated by the model after shifting and re-scaling them to lie on the unit interval. The red indicates hierarchy values passed through a logistic function to remove the nonlinearity introduced by the logit link function in the GLM fitting procedure. The monotonicity of this transformation preserves the order of the cortical regions and therefore does not affect the Spearman rank correlations reported in the main text. **(d)** Myelin map values for 89 cortical areas.

**Extended Data Figure 3:**
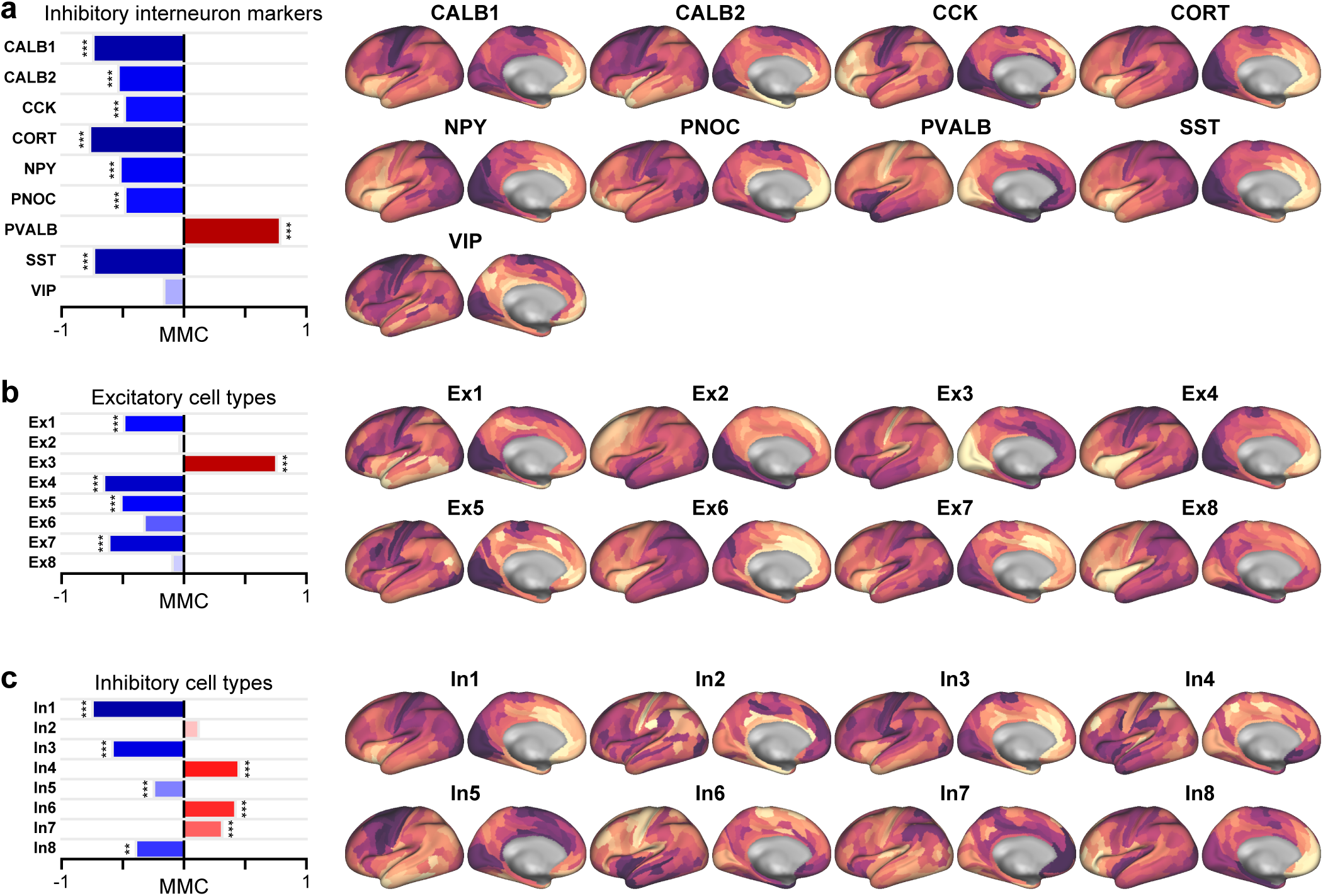
Expression maps and MMCs for genes that code for markers of distinct inhibitory interneuron cell types, and for weighted profiles characteristic of distinct neuronal cell types derived from single-cell RNA sequencing of human cortical neurons. **(a)** Markers for inhibitory interneuron cell types. **(b)** Weighted gene sets for excitatory neuronal cell types, derived from single-cell RNA sequencing. **(c)** Weighted gene sets for inhibitory neuronal cell types, derived from single-cell RNA sequencing.

**Extended Data Figure 4:**
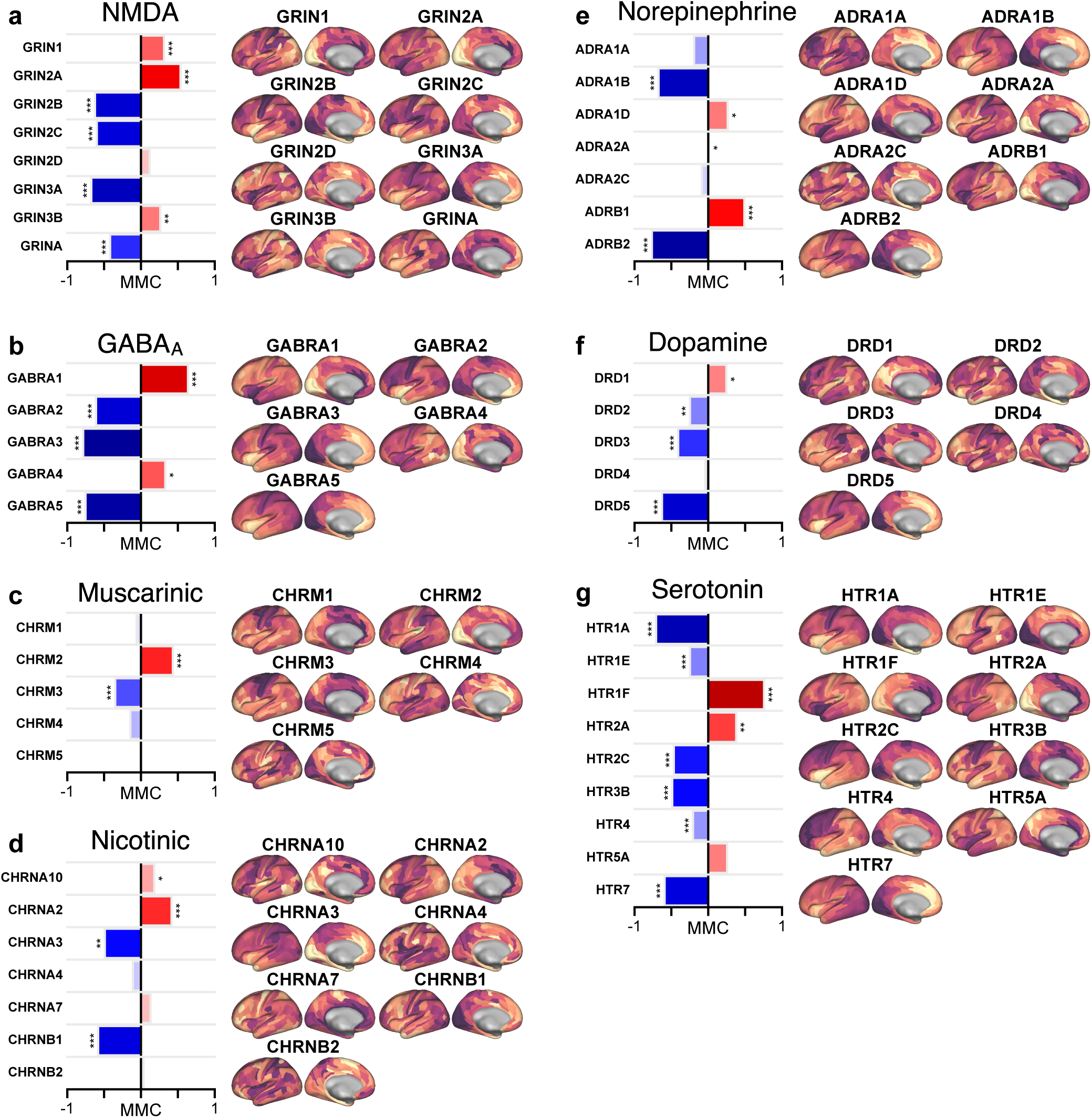
Expression maps and MMCs for genes coding for synaptic receptor subunits and neuromodulator receptors. **(a)** NMDA receptor subunits. **(b)** GABA_A_ receptor subunits. **(c)** Muscarinic acetylcholine receptors (CHRM). **(d)** Nicotinic acetylcholine receptors (CHRN). **(e)** Norepinephrine receptors (ADR). **(f)** Dopamine receptors (DRD). **(g)** Serotonin receptors (HTR).

**Extended Data Figure 5:**
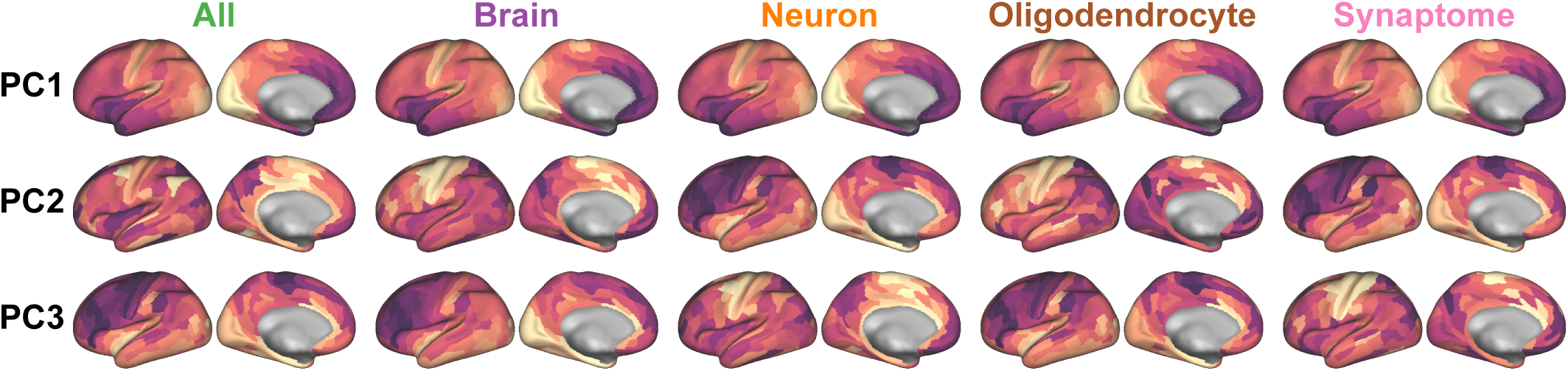
Principal component analysis (PCA) reveals that the dominant axis of gene expression (i.e., the first principal component, PC1) is conserved across categorical gene sets. PC1, which aligns with the myelin map, separates sensory areas (e.g., primary visual, somatosensory, and auditory cortical areas), from association areas. For each gene set, the secondary and tertiary main of gene expression (i.e., PC2 and PC3) tend to fractionate sensorimotor cortical areas by modality, separating early visual cortex and somatomotor cortex. Rows correspond to the first three PCs, respectively, across gene sets.

**Extended Data Figure 6:**
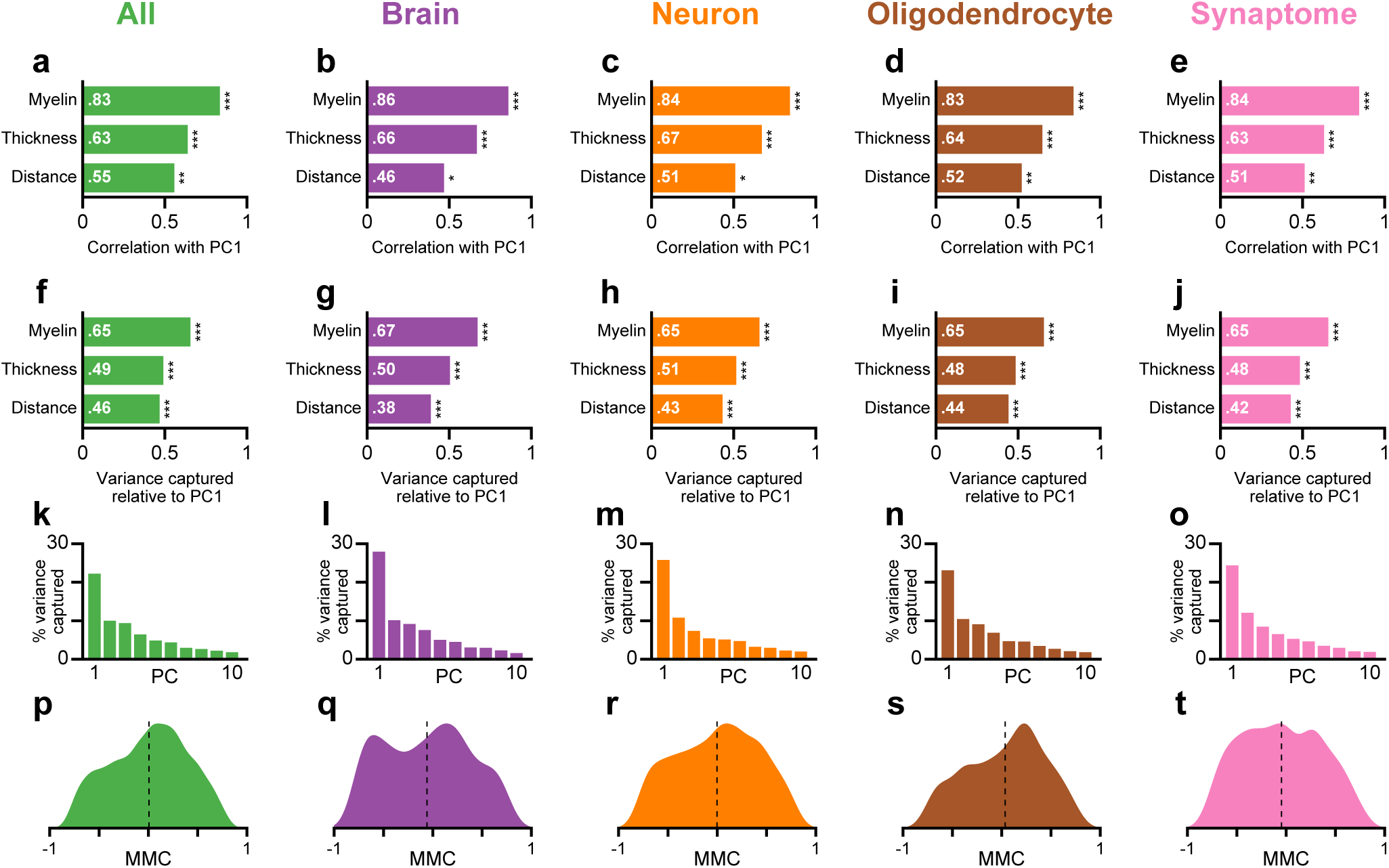
The dominant axis of gene expression is better captured by the myelin map than by two alternative candidate proxies: cortical thickness and distance from primary visual cortex (V1). **(a–e)** For five gene sets, the Spearman rank correlation between the first principal component (PC1) and the cortical myelin map, the map of cortical thickness, and the map of geodesic distance from primary visual cortex. For each gene set, PC1 is more strongly correlated with the myelin map than with the two other candidate maps. **(f–j)** For five gene sets, the amount of gene expression variance captured, relative to PC1, for the three candidate maps. For each gene set, the myelin map captures more gene expression variance than do other two maps. **(k–o)** Percentage of gene expression variance captured by the top 10 PCs, out of 179 total PCs (due to 180 cortical areas in our parcellation). For all five gene sets, PC1 captures between 22% and 28% of the variance, which is more than twice the amount captured by PC2. **(p–t)** Distribution of myelin map correlations (MMCs) across genes for the five gene sets. Dashed lines mark the mean of the distribution. For all five gene sets, the distributions are broad, containing large fractions of strong positive and negative MMCs, and centered near zero, with a range of means (*−*0.06, +0.04).

**Extended Data Figure 7:**
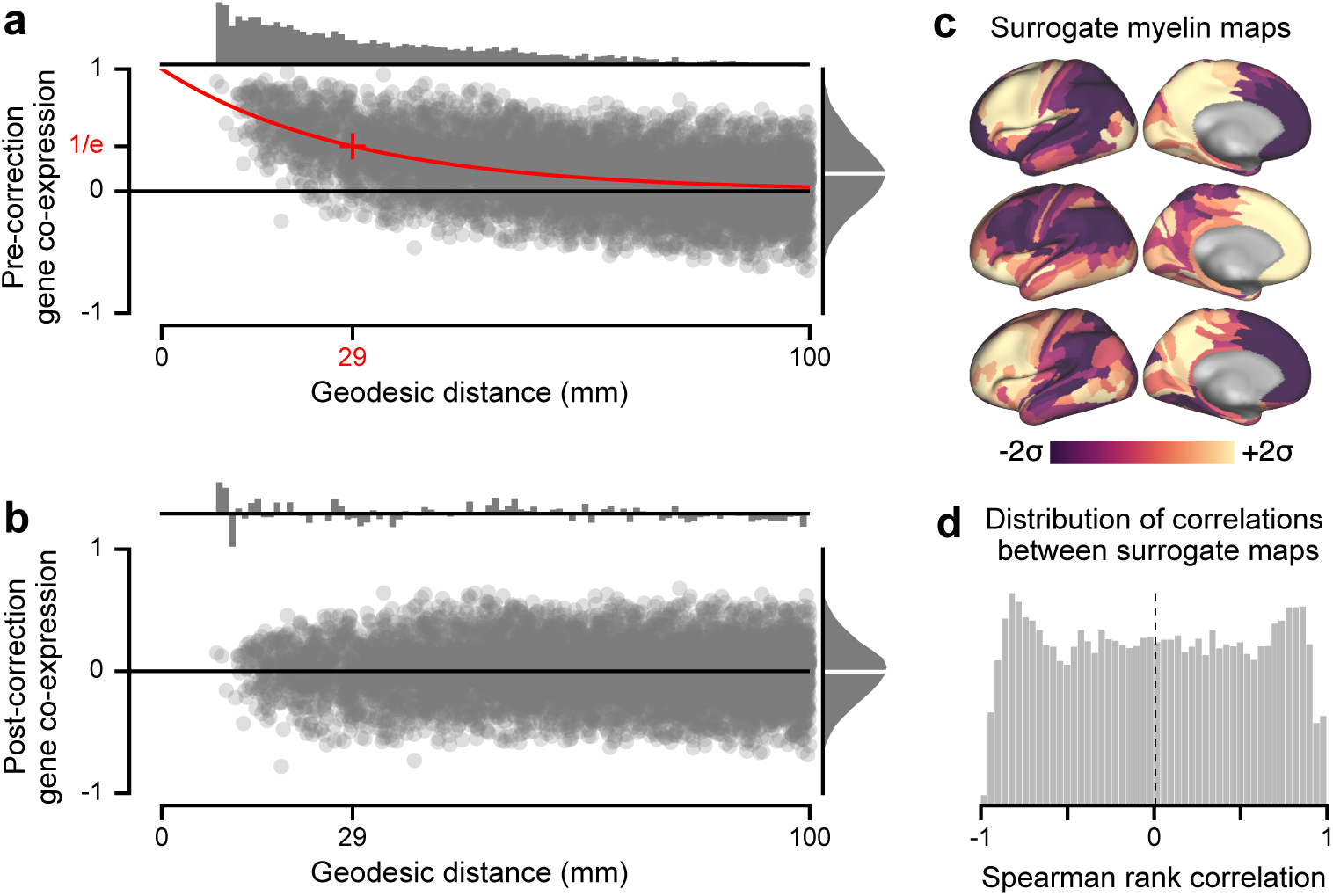
Autocorrelation structure in gene expression and myelin maps. **(a)** Spatial autocorrelation structure in the parcellated cortical gene expression data is well-approximated by a decaying exponential. Gene co-expression is defined as the pair- wise Spearman rank correlation between cortical parcels’ gene expression values, here for the brain-specific gene set. Proximal cortical parcels exhibit more similar gene expression values compared to distal parcels. All pairs of parcels with geodesic distance less than 100 mm were used to fit the characteristic scale of spatial autocorrelation, illustrated in red (i.e., exp(−*d/d*_0_)), where *d* is geodesic distance and *d*_0_ = 29 mm. Each data point corresponds to the co-expression of a pair of cortical parcels. *Top*: Mean co-expression value as a function of geodesic distance bin. **(b)** Gene co-expression values after correcting for spatial autocorrelation structure by subtraction of the fitted exponential decay. After correction, the mean co-expression value is near zero across all geodesic distance bins. **(c)** Example randomized surrogate maps with spatial autocorrelation structure matched to the cortical myelin map (see Methods). Autocorrelation structure-preserving surrogate myelin maps are used for nonparametric calculation of statistical significance for PCA results in Figs. 5 and 6. **(d)** Distribution of pairwise Spearman rank correlations between pairs of surrogate myelin maps.

